# Toxic Effects of Crude Extracts from Chinese Herbal Toothpastes on *Solenopsis invicta* (Hymenoptera: Formicidae)

**DOI:** 10.1101/648139

**Authors:** Qian Xiao, Babar Hassan, Junzhao Luo, Yijuan Xu

## Abstract

Red imported fire ant is an invasive species with the characteristics of quick dispersion, strong ferocity and aggression. For a long time, chemical control has been used as a main means of prevention and control of this pest, but the long-term use of high toxic pesticides will lead to a serious impact on the environment and non-target organisms. The toxicity of Chinese herbal toothpaste and its crude extracts against red imported fire ant was evaluated in the laboratory bioassay. The results showed that the traditional Chinese medicine toothpaste with watermelon cream has significant toxicity to red imported fire ants. The mortality reached more than 80% after 48h treatment, and the mortality was 100% after 72h treatment. Among the fractions separated from watermelon cream after the extraction of toothpaste, the ethyl acetate fraction has higher activity, and the mortality was more than 80% when the concentration was 1% after 72h. Our results suggest that Chinese herbal toothpaste is toxic to insects and have the potential for bait application in pest control.

## Introduction

*Solenopsis invicta* Buren, the red imported fire ant, is an important invasive insect with the characteristics of quick dispersion, strong ferocity and aggression. It causes severe damage to agriculture and poultry production, ecosystem and human health [1]. At present, chemical control is the most widely used method to control red imported fire ants. Several insecticides have been registered and used against this insect pest [2]. These insecticides are mainly used by flooding the mounds with water, as a powder and poisonous baits [3]. Carbaryl, diazinon, chlorpyrifos and some other insecticides are used for flooding the mounds. The main components of powder are cypermethrin and fluronitrile. Similarly, several slow releasing insecticides and insect growth regulators are usually used as poisonous bait [3].

Although chemical control works well, but the major concern in using these insecticides is, their negative effect on environment and these are also harmful to non-target organisms [4]. Moreover, sustained insect resistance against synthetic insecticides has also invigorated the development of unconventional insect pest management practices. Therefore, search for new insecticide with different mode action, low mammalian toxicity, having less impact on biodiversity and environment is major theme of pest control. Several plants derived chemicals have been used as a safer and green pesticides against red imported fire ants. For example, two terpenoids isolated from leaves of American beautyberry (*Callicarpa americana* L.) and Japanese beautyberry (*Callicarpa japonica* Thunb.), and essential oil from *Cupressus nootkatensis* (D. Don) showed repellant and toxic effects against *S. invicta, S. richteri* Forel, and a hybrid of these two species at very low concentrations [5, 6]. Both clove powder and clove oil have shown promising results in against the red imported fire ant [7]. Natural sweetener, erythritol, has a good toxic effect on red imported fire ants, and significantly affected the foraging behavior and necrophoric behavior of the fire ants [8]. The food additives such as ethyl anthranilate and butyl anthranilate, can prevent fire ant nesting in pots [9].

The active ingredients of Chinese medicine are extracted from Chinese herbal plants, such as *radix zanthoxyli* (dried root of *Zanthoxylum nitidum* (Roxb.) D.C.), *Morus alba* L., *Rhizoma rehmanniae* dihuang *Artemisia vulgaris* L., *Coptis chinensis* Franch, *Panax pseudoginseng* (Burk.), *Taraxacum* spp. and many others plants [10, 11]. Previously, several studies showed that some active ingredients in traditional Chinese medicine have toxic effects against several microorganisms and pests [12]. For example, the extracts of *Radix aconitum* L., *Gelsmium elegans* Benth, *Impatiens balsamina* L. and *Polygala tenuifolia* Willdenow can inhibit the growth of hypha of tomato necrotrophic fungus, *Botrytis cinerea*, and the inhibition rate increases as the concentration of the extracts increases [13]. Three traditional Chinese medicine of *Cinnamonum cassia, Eugenia caryophyllata* and *Pogostemon cablin* have strong insecticidal activity against house dust mites [14]. Similarly, the constituents of patchouli oil extracted from *Pogostemon cablin* (Blanco) BENTH has great acaricidal activity against house dust mite, *Dermatophagoides farinae* (Wu et al., 2012).

To our knowledge, the potential of Chinese herbs or herbal medicine as a control agent of fire ants have not been tested, previously. Therefore, in this study, we wanted to test different Chinese herbal toothpastes with good control effect on red imported fire ants and low mammalian toxicity to achieve sustainable development in less toxic pesticides and effective management of red imported fire ants.

## Materials and methods

### Insect

Red imported fire ant, *S. invicta*, were collected from Tianhe District Wisdom Park, Guangzhou, Guangdong province, China. Spades were used to dig the nest of ants, and these were moved quickly into plastic barrel. Before digging, plastic barrel walls, spade handles, rubber gloves and long rainwater boots were coated with Fluon to prevent red fire ants from climbing. After collection, the inner and outer walls of the barrel were coated with Fluon again to avoid escape of ants [15].

The ant colonies were raised in plastic boxes (40cm×26cm×10cm), with Fluon solution coated in the inner wall to prevent escape of insects. The indoor feeding temperature was 25 ± 1 °C and humidity was 60 – 70%. We placed petri dishes (9 cm dia) with wet gypsum as an artificial nest for ant colonies in each plastic box. A test tube (25 by 200 mm) that was partially filled with water and plugged with cotton was used as a water source. The fire ants were given a 10% solution of honey mixed with water (50 ml) weekly. Frozen locust, *Locusta migratoria manilensis*, were used as a food source.

### Reagent

Bright blue dye (Zhengxing Food Additive Co., Ltd., Zhengzhou, Henan Province), PBS buffer, ethanol (Shengxinyuan Weiye Trade Co., Ltd., Hebei District, Tianjin), ethyl acetate (Shengxinyuan Weiye Trade Co., Ltd., Hebei District, Tianjin), sucrose (Taigu Sugar Industry China Co., Ltd., Guangzhou, Guangdong province), Yunhan Watermelon cream toothpaste (Guilin Yunhan Daily Chemical Co., Ltd.), Pianzaihuang toothpaste (Zhangzhou Pianzaihuang Shanghai Jiahua Oral Care Co., Ltd.), the details of ingredients and their manufacturers of twelve traditional Chinese medicine toothpaste is presented in Table 1.

### Comparison of toxicity of different traditional Chinese medicine toothpaste to red imported fire ants

In order to compare the toxicity of different traditional Chinese medicine toothpaste to red imported fire ant, about 30 workers were selected randomly and put into plastic bowl (3cm radius, 6cm high) after 12 hours of starvation. The plastic bowl was coated with Fluon to prevent ants from escaping. 10% of each toothpaste was prepared using distilled water as a solvent. A 2-ml centrifuge tube plugged with cotton with toothpaste solution was then introduced into the plastic bowl. Distilled water and 10% sucrose solution were used as control treatment in the experiment and each treatment was repeated thrice. Mortality of workers was recorded after every 24 hours until three days and percentage mortality were calculated.

### Confirmation of toothpaste intake by ants

In order to verify whether the workers have consumed the solution of traditional Chinese medicine toothpaste, the bright blue dye was mixed with the 0.01M PBS to produce 0.5% solution of dye. A total of 50µl dye solution was added to each Chinese traditional medicine toothpaste solution (10%) of watermelon cream and pinzaihuang. The mixture was fed to the ants in the same way as in toxicity test. Distilled water and 10% sucrose solution were used as a control treatment. Each treatment was repeated three times. After 24 hours of feeding, the abdomens of the workers were squeezed to observe the presence of dye in the abdomen and percentage of insects have dye in the abdomen were calculated.

### Toxic effects of Watermelon Cream toothpaste against Red Imported Fire ants

In screening test, we found that Watermelon Cream toothpaste is most toxic to the workers. In order to compare the toxic effects of different concentrations of watermelon cream toothpaste solution against red imported fire ants, the toothpaste was mixed with distilled water at five different gradient concentrations: 0.5%, 1%, 5%, 10% and 20%. Toothpaste solution was added into centrifuge tubes (2 ml) and was plugged with small amount of cotton. Thirty workers were randomly selected after 12 hours starvation and were released in the plastic bowls. Tubes containing toothpaste solution were placed in the plastic bowls for feeding of ants. Distilled water and 10% sucrose solution were used as control treatment. Each treatment was repeated three times. Number of dead ants was recorded after every 24 hours until three days.

### Extraction of Yunhan Watermelon Cream toothpaste

In order to identify the toxic active components of toothpaste to red imported fire ant, we soaked 1000.00 g Yunhan watermelon cream toothpaste in 380 ml ethanol (purity ≥ 99.7%) and solution was stirred which lasts for 4 hours. This was repeated three times at room temperature. White precipitates (S-1) of 656.3 g were obtained by drying filterare residue.

Ethanol extract of toothpaste was obtained when filtrate was concentrated by rotating evaporation (50 °C). We added water to the ethanol extract and extracted it with ethyl acetate. The water layer and ethyl acetate layer were concentrated by rotating evaporation to obtain the water extract (S-2) of 202.8 g and the ethyl acetate extract (S-3) of 114.5 g (Fig. S1).

**Fig. S1** Extraction steps of Yunhan watermelon cream toothpaste.

### Toxic effects of extracts from Yunhan Watermelon Cream on Red Imported Fire ants

In order to compare the toxic activity of each component of the Yunhan Watermelon Cream toothpaste to the red imported fire ant, the three extracts of toothpaste, S-1, S-2, and S-3, were diluted into 1% and 10% concentration in their respective solvents and were added in 2mL centrifuge tubes. The tubes were plugged with cotton as describes earlier. Thirty workers were randomly selected, after 12 hours of starvation and were released in plastic bowls. Tubes containing toothpaste extracts (S1-S3) were placed in plastic bowls to feed red fire ants. Distilled water and 10% sucrose solution were used as control treatment and each treatment was repeated three times. Number of dead red fire imported ants was recorded after every 24 hours until three days and percentage mortality were calculated.

### Selective preference of Red Imported Fire Ant to S-3 extract

In order to assess whether toothpaste extract S-3 affect foraging preference of fire ants, 700 workers were randomly selected into rectangular plastic box (17cm × 11.5cm × 5.5cm) after 12 hours of starvation. The walls of plastic box were coated with Fluon to prevent ants from escaping. We took two 1.5mL centrifuge tubes, one containing 10% sucrose solution, the other containing the 0.5% toothpaste extract solution (S-3 extract + 10% sucrose). Small holes about the diameter of 1 mm were made near the mouth of tubes so that the solution does not flow out when inverted but the ants can suck it. After labeling, these were weighed using analytical balance. Both centrifuge tubes were placed upside down on the two ends of the plastic box respectively. After 24 hours, the centrifuge tubes were taken out to weigh again. The actual amount of food consumed by red fire ants over the past 24 hours was obtained through the weight difference of the centrifuge tube. The control group was made by same material and environment without fire-imported ants. The amount of food consumed by ants of three different concentrations (0.5%, 1% and 2%) of S-3 and sucrose solution was calculated.

### Data Analysis

All percentage mortality data were transformed using the arcsine transformation to meet the assumptions of normality. Analysis of variance (ANOVA) followed by Tukey’s test for multiple was used for analysis of obtained data at 5% level of significance using SPSS 22.0. Binary choice tests to determine feeding preferences of ants for watermelon extracts were analyzed using a paired t-test comparison.

## Results

### Comparison of toxicity of different traditional Chinese medicine toothpaste to red imported fire ants

Results of toxicity test showed that there was no significant difference between different toothpaste treatments in term of mortality of ants in the first 24 hours (*P* = 0.105). But there was significant difference in the mortality of red imported fire ants after 48 hours of treatment with twelve traditional Chinese medicine toothpaste and control treatment (*F* = 3.30, *df*_*1*_ = 13, *df*_*2*_ = 28, *P* = 0.004). Among them, the mortality of watermelon cream and Pinzaihuang was more than 80%, which was significantly higher than that of other toothpaste. The mortality of ten toothpaste was less than 70%, including four type of toothpaste of *Radix zanthoxyli*, three type of toothpaste of Saky, three type of toothpaste for Heimei, Yunnan Baiyao, and Pudilan. In the control group, the mortality ants on distilled water and sucrose was less than 30 and 10%, respectively (Fig. 1).

**Fig. 1.**
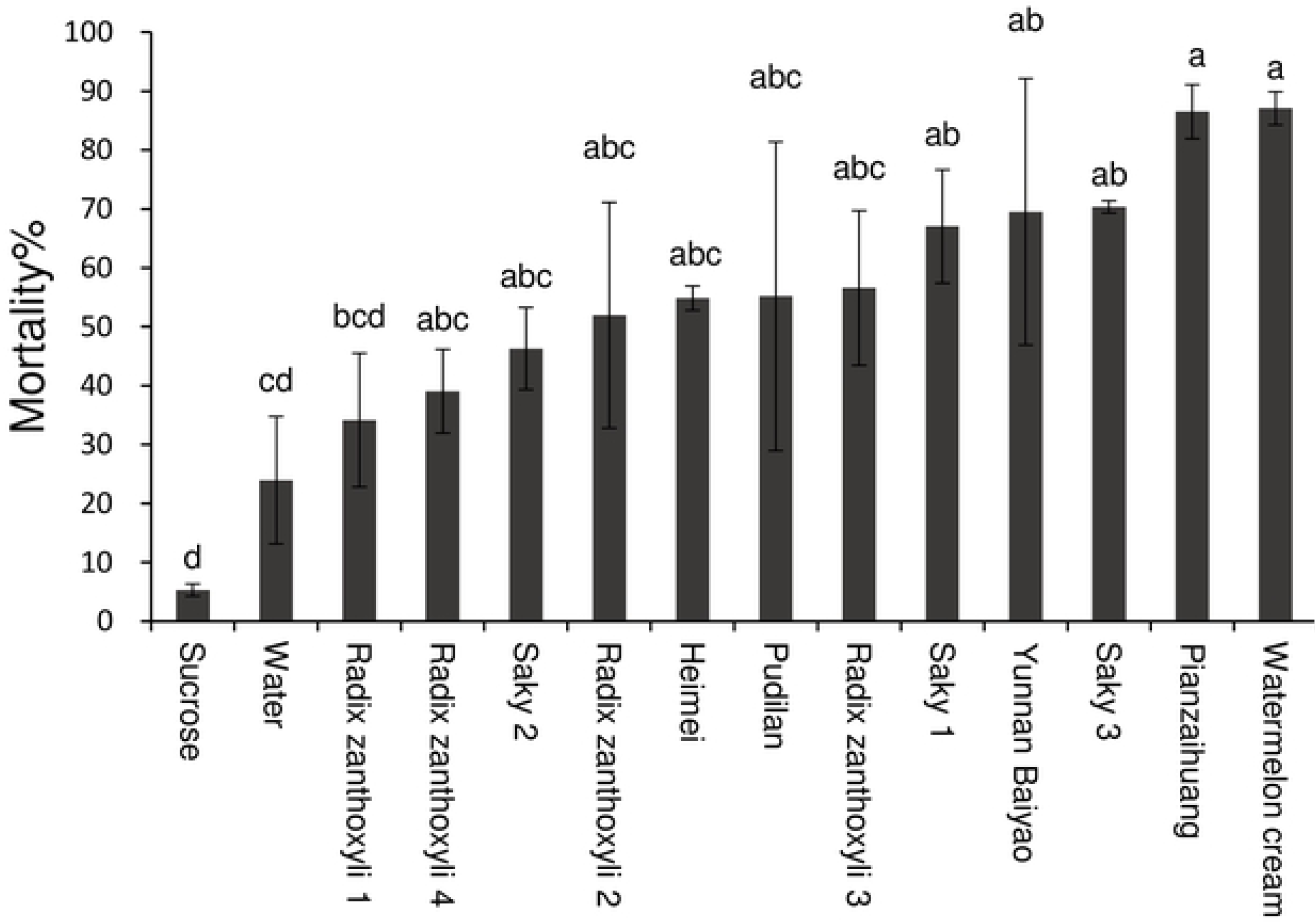
Comparison of mortality (mean ± standard error) of red imported fire ants after feeding on traditional Chinese medicine toothpastes for 48 hours. Bars with the same letter do not significantly differ (Tukey’s test) at the 0.05

#### significance threshold

There was also a significant difference in mortality of red imported fire ants after 72 hours treatment after feeding on twelve type of traditional Chinese medicine toothpaste (*F* = 8.484, *df*_*1*_ = 13, *df*_*2*_ = 28, *P* < 0.0001). Among them, the mortality rates of several traditional Chinese medicine toothpaste solutions, including *Radix zanthoxyli* 4, Saky 1, Saky 3, Yunnan Baiyao, Pudilan, were more than 90%, while the death rates of watermelon cream and Pianzaihuang were 100%. The mortality rates of *Radix zanthoxyli* 1, *Radix zanthoxyli* 2, Saky 2 and Heimei were less than 80%. In the control group, the mortality of ants after feeding on distilled water and sucrose was was less than 39.97 and 20 %, respectively (Fig. 2).

**Fig. 2.**
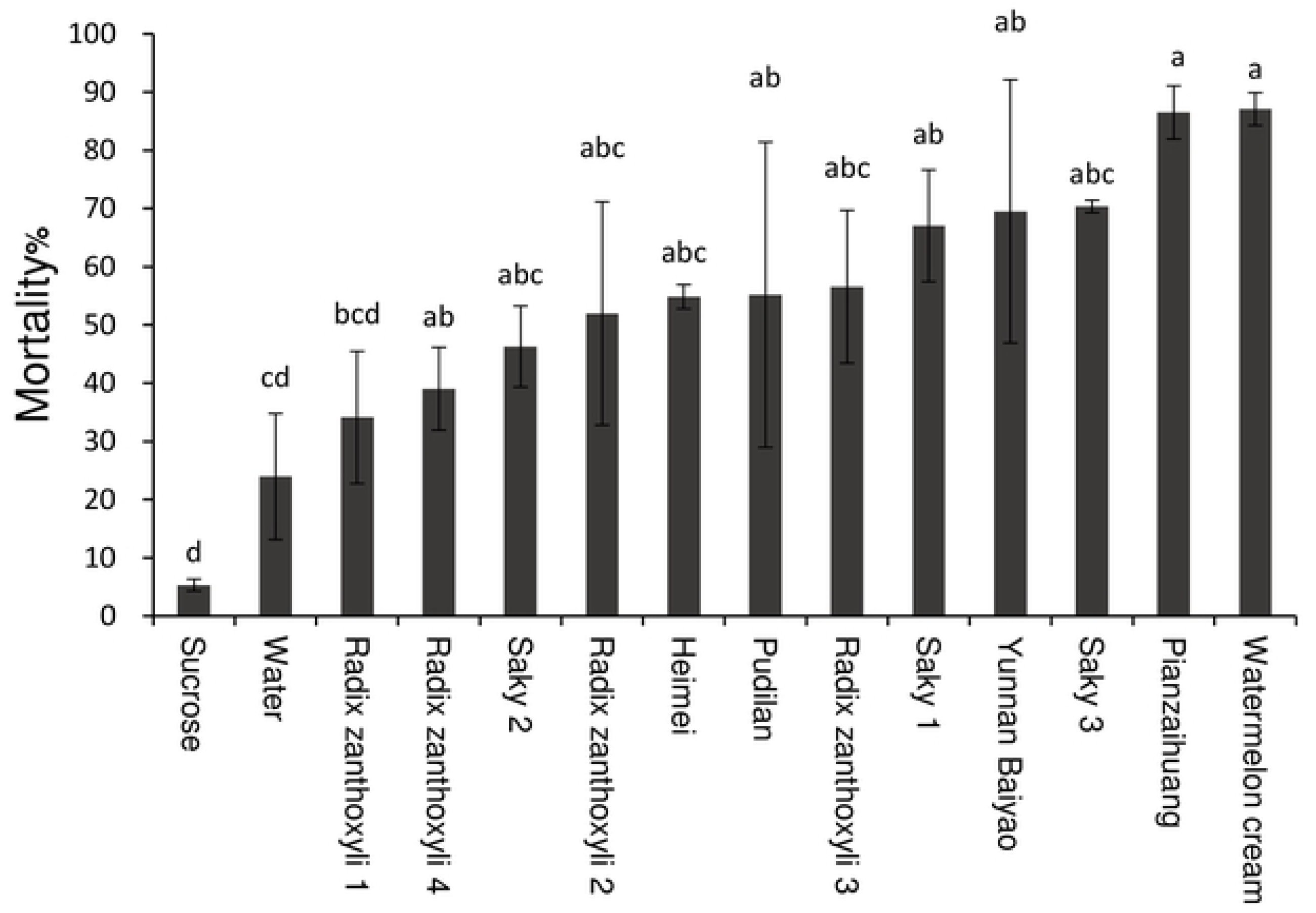
Comparison of mortality (mean ± standard error) of red imported fire ants after feeding on traditional Chinese medicine toothpaste for 72 hours. Bars with the same letter do not significantly differ (Tukey’s test) at the 0.05 significance threshold.

### Confirmation of toothpaste intake by ants

The toothpaste solutions of watermelon cream and Pianzaihuang, which had the highest mortality for ants in the selection test, were selected for confirmation of food (toothpastes) intake by ants and dyeing rate was calculated. After different treatments, the dyeing rates of red imported fire ants were significantly different (*F*=584.9778, *df*_*1*_=3, *df*_*2*_=8, *P*<0.0001). The staining rate of watermelon cream was close to 80%, but only one red fire ant was stained in one repeated experiment in Pianzaihuang. In the control group, the dyeing rate of sucrose and distilled water was 99% and 100% (Fig. 3).

**Fig. 3.**
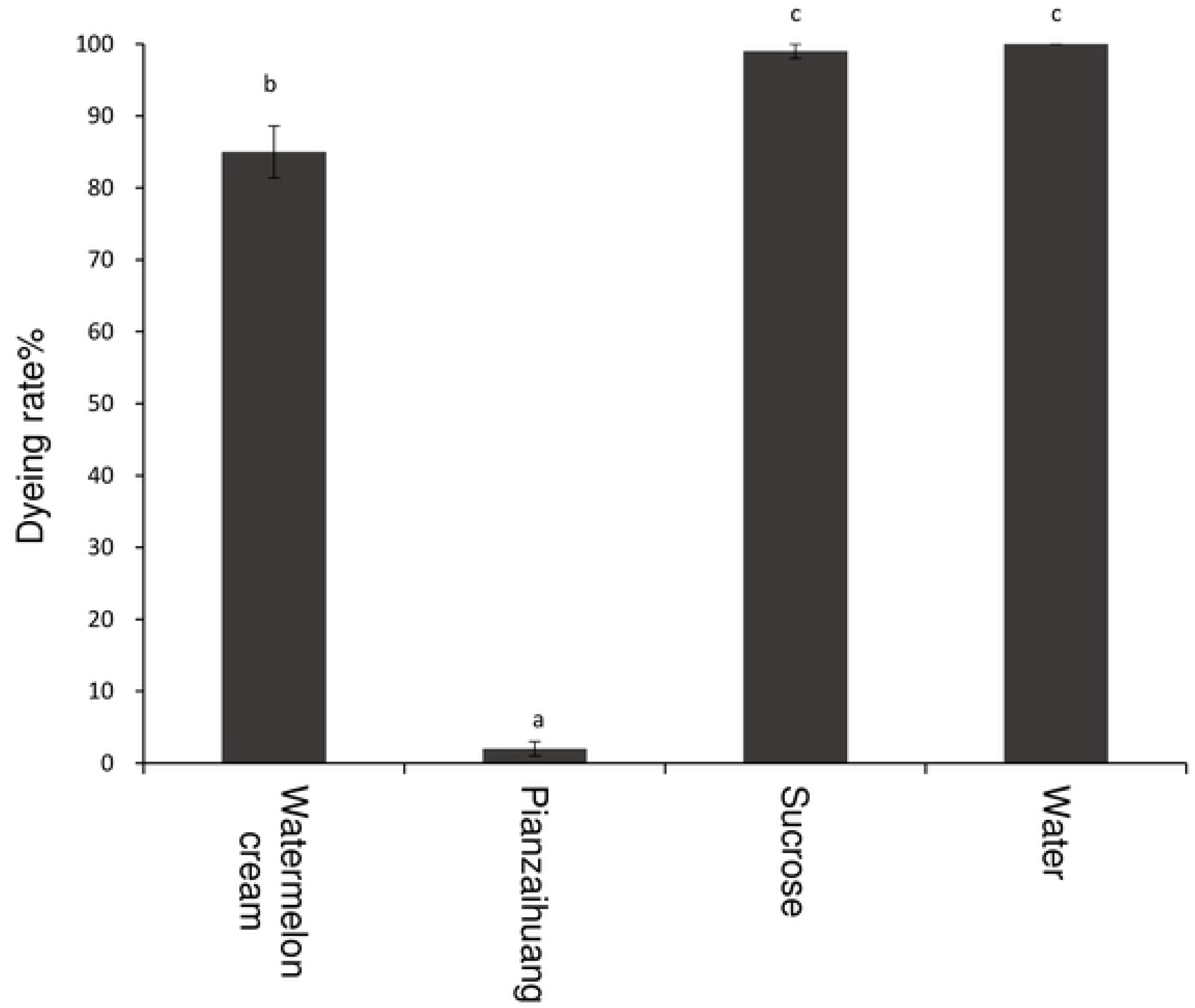
comparison of dyeing rate (mean ± standard error) of red imported fire ants after feeding of 24 hours on two Chinese medicine toothpastes. Bars with the same letter do not significantly differ (Tukey’s test) at the 0.05 significance threshold.

### Toxicity of watermelon cream toothpaste against Red Imported Fire Ants

The toxic effects of different concentrations of watermelon cream toothpaste solution on red fire ants were significantly different (*F* = 60.036, *df*_*1*_ = 6, *df*_*2*_ = 54, *P* < 0.0001). The mortality of red fire ants increased over time (*F* = 17.301, *df*_*1*_ = 4, *df*_*2*_ = 54, *P* < 0.0001). The toxic effect was remarkable at the concentration of 10% and 20%. The mortality of ants at the concentration of 0.5%, 1% and 5% were close to that of water. The mortality rates were low, and the toxic effects were not significant. The mortality at concentration of 1% was less than that in water (Fig. 4).

**Fig. 4.**
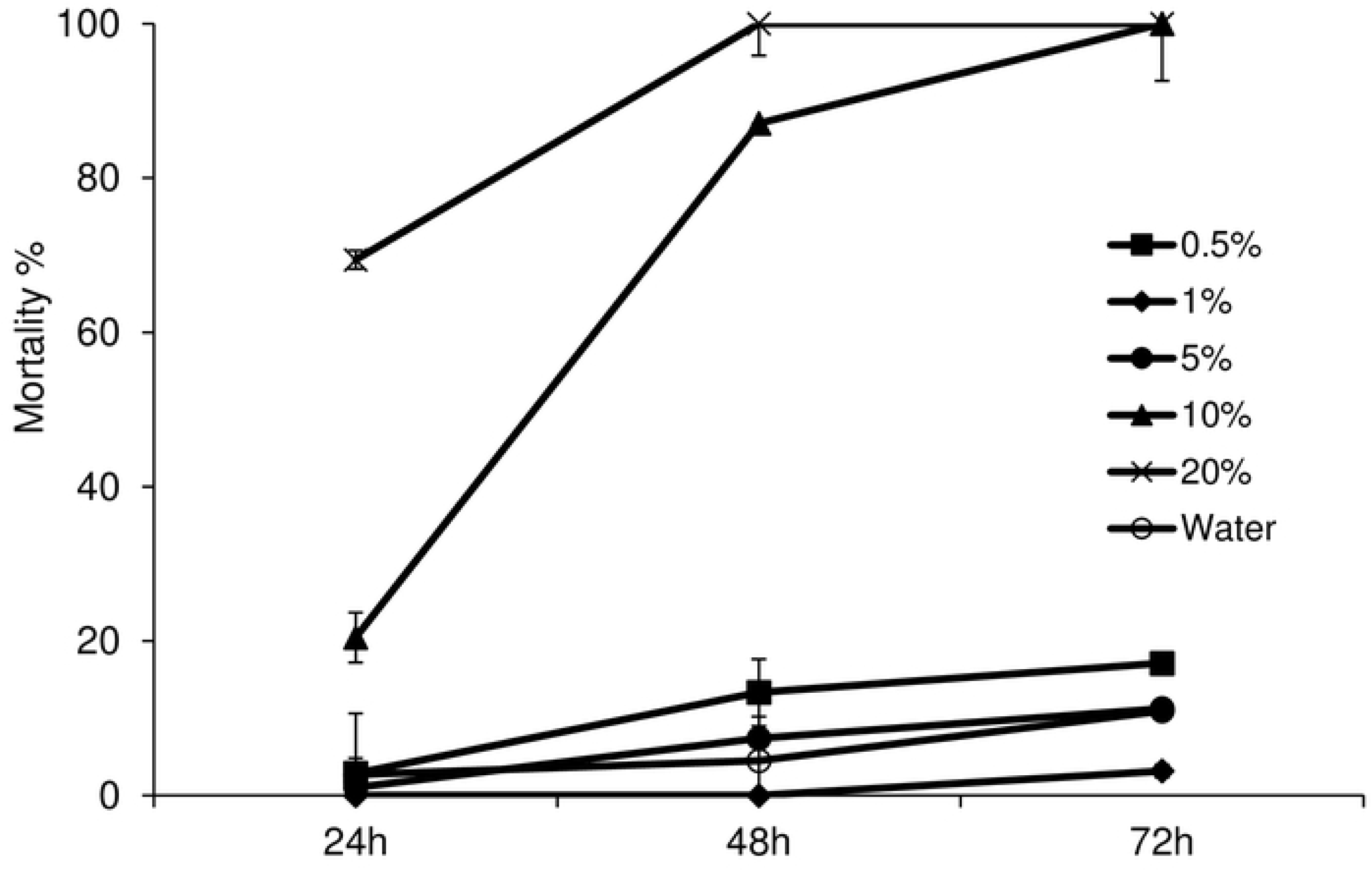
Comparison of mortality (mean ± standard error) of red imported fire ants after feeding on different concentrations of watermelon cream solution.

### Toxic effects of extracts from Watermelon Cream on Red Imported Fire ants

The toxic activities of different components isolated from Watermelon Cream toothpaste against red imported fire ant at concentration of 10% were significantly different (*F* = 46.633, *df*_*1*_ = 4, *df*_*2*_ = 38, *P* < 0.0001). The mortality of red fire ants increased over time (*F* = 83.307, *df*_*1*_ = 2, *df*_*2*_ = 38, *P* < 0.0001). Among them, the solution of watermelon cream toothpaste at concentration of 10% had significant toxic effect on ants. The mortality of ants after feeding on S-2 and S-3 at 10% concentration were higher than that of 10% watermelon cream toothpaste solution (without extraction) after 24 hours and 48 hours. Moreover, within 48 hours, all ants were dead (100% mortality). The mortality of S-1 at 10% concentration was significantly less than that of toothpaste solution (Fig. 5).

**Fig. 5.**
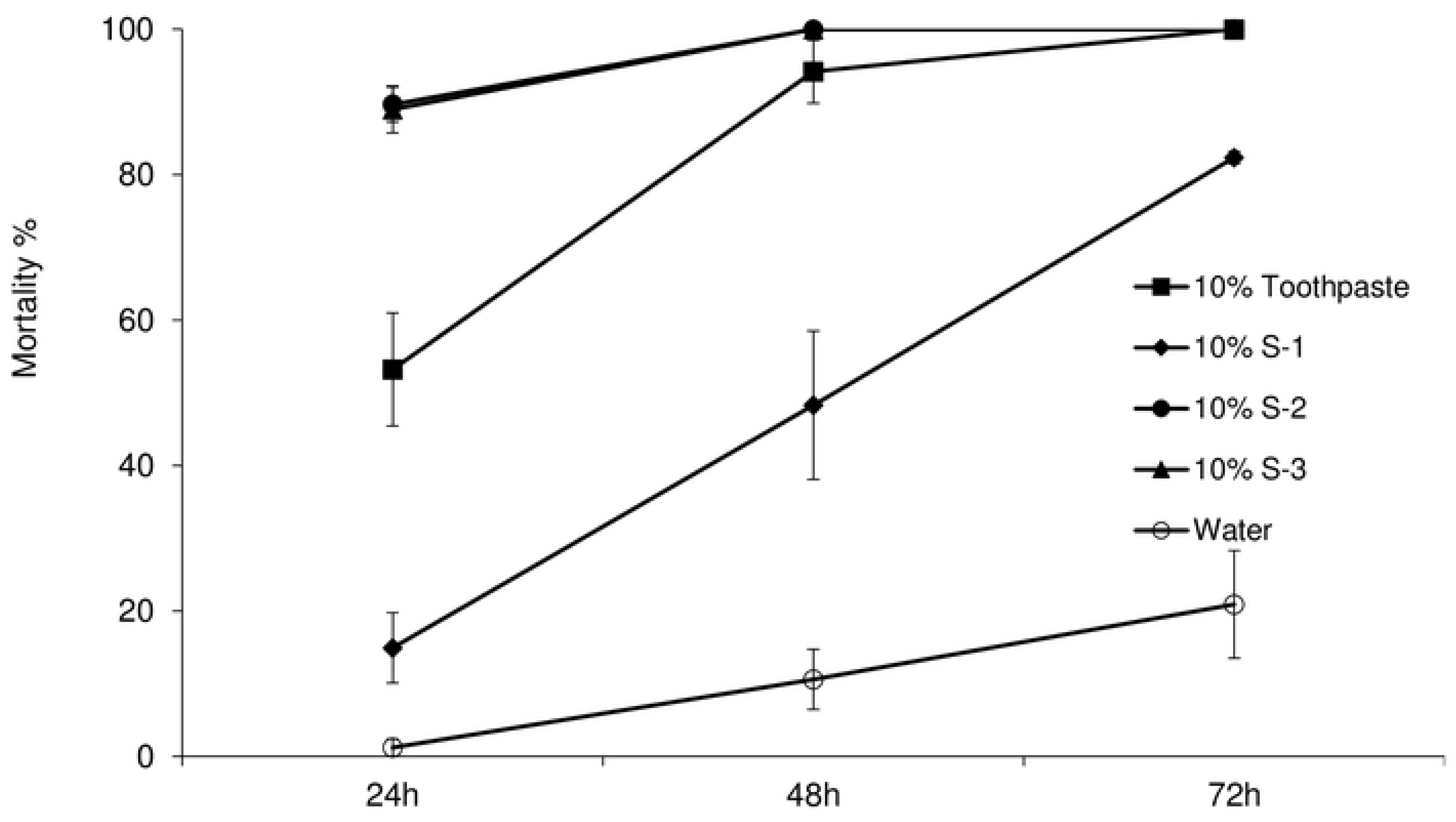
Comparison of mortality (mean ± standard error) of red imported fire ants after feeding on different parts of watermelon cream at 10% concentration.

Similarly, the toxic effects of different components of watermelon cream toothpaste on red fire ants were significantly different at 1% concentration (*F* = 77.713, *df*_*1*_ = 4, *df*_*2*_ = 38, *P* < 0.0001). With the passage of time, the mortality of ants increased (*F* = 77.713, *df*_*1*_ = 4, *df*_*2*_ = 38, *P* < 0.0001). After 72 hours, only the mortality of ants after feeding on S-3 at the concentration of 1% was more than 80%, while the mortality on 1% S-1 and S-2 were less than 80%. The mortality in water treatment was less than 30% (Fig. 6).

**Fig. 6.**
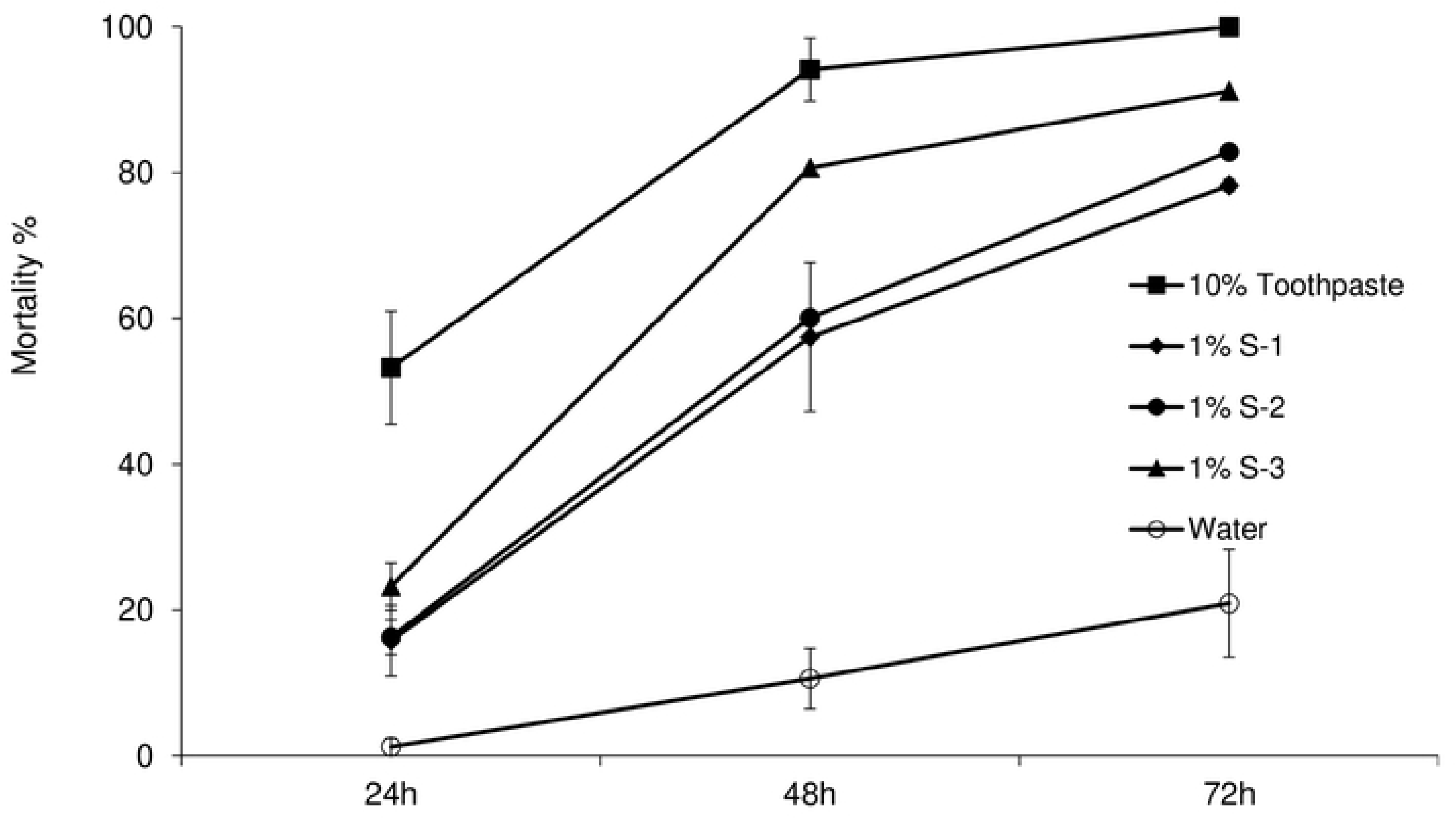
Comparison of mortality (mean ± standard error) of red fire imported ants after feeding on different parts of watermelon cream at 1% concentration.

### Selective preference of Red Imported Fire Ant to S-3 extract

The results showed that there was no significant difference in the intake amount of S-3 at the concentration of 0.5% or 2% of ants compared with sucrose solution (0.5%: *t* = 1.779, *df* = 8, *P* = 0.113; 2%: *t* = 1.122, *df* = 8, *P* = 0.294). However, the intake of 1% S-3 was much more than that of the control (*t* = 3.26, *df* = 8, *P* = 0.015) (Fig. 7).

**Fig. 7.**
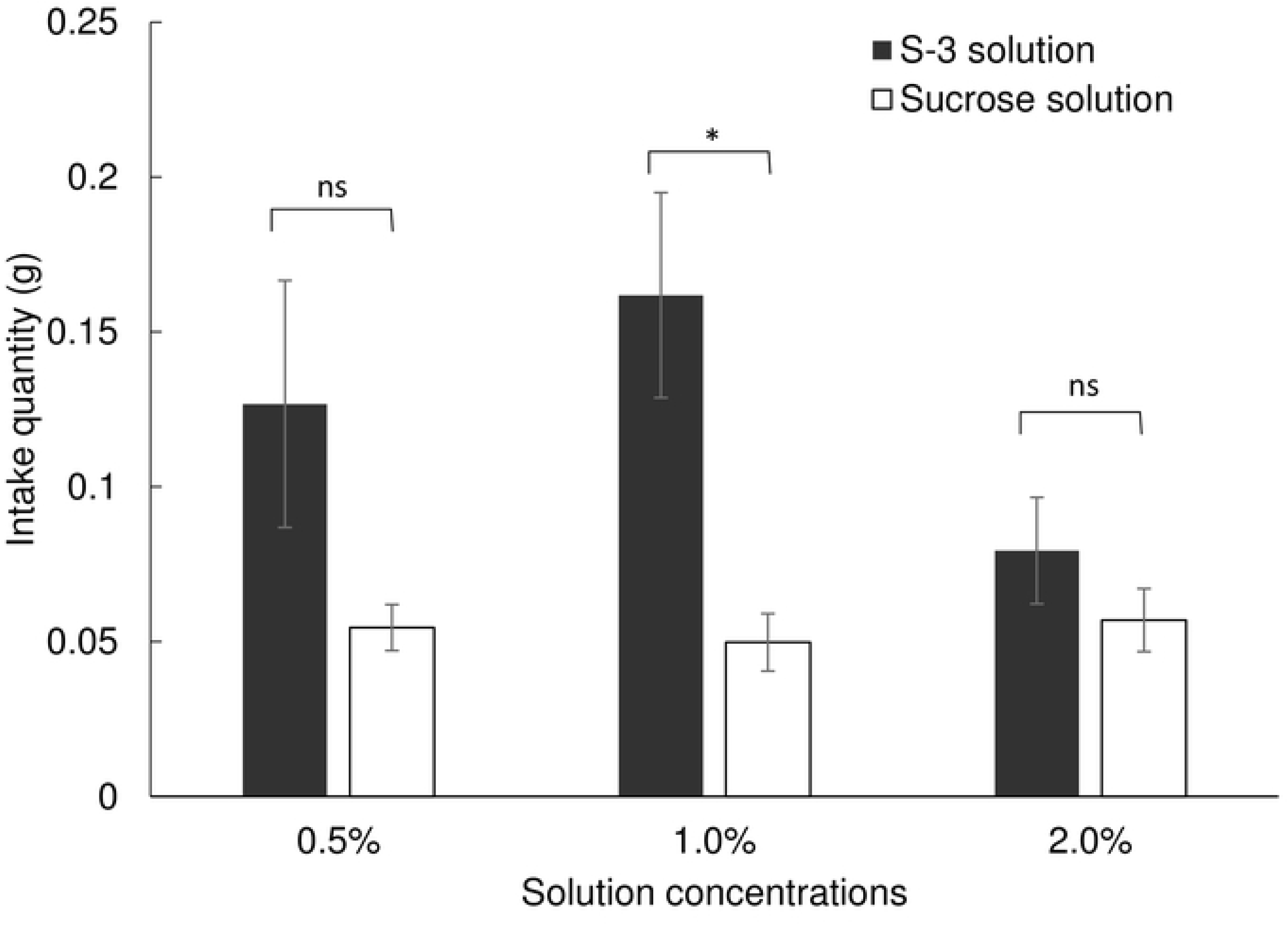
Comparison of intake amount by red imported fire ant between S-3 solution (mean ± standard error) and sucrose solution. * or ns above the bars indicates statistically significant (*P* < 0.05) or not significant (*P* > 0.05) differences between S-3 solution and sucrose solution treatments (Paired t test).

## Discussion

In this study, Chinese medicine toothpaste of Yunhan Watermelon Cream had strong toxicity to red imported fire ants. The mortality of workers fed on extract S-3 solution at the concentration of 1% was over 80% after 72 hours, which indicates that the traditional Chinese medicine toothpaste may be significantly more toxic than food additives such as erythritol and glycine [16]. Acute toxicity test and long-term toxicity test showed that Watermelon Cream toothpaste is safe for daily use and clinical application. Therefore, it is possible to screen out toxic active ingredients from Watermelon Cream toothpaste for fire ants, which are environmentally friendly and harmless to human.

In the experiment on toxicity of different concentrations of Yunhan Watermelon Cream toothpaste solution to red imported fire ant, we found that the high concentration of watermelon cream solution was toxic to workers, and the low concentration of watermelon cream solution did not affect the survival of ants significantly and even provide nutrients for them. The active ingredients of this toothpaste are *Panax notoginseng, Coptis chinensis* that has been proven to be of great benefits to human health. However, previous studies showed that flower buds extract from *p. notoginseng* containing triterpene saponins that were toxic to larvae and pupae of *Aedes albopictus*c [17]. This plant contains ginsenosides and notoginsenoside as major constituents, which are toxic to insects and several microorganisms. The main chemical constituents of *C. chinesis* are quinoline alkaloids, including berberine, coptisine, palmatine, and jatrorrhizine. Naturally, occurring alkaloids in many plants are as non-toxic N-oxides. However, as soon as they reach the often-alkaline digestive tracts of some insect herbivores, they are quickly reduced and forms toxic, uncharged, hydrophobic tertiary alkaloids and cause mortality of insect [18, 19]. Isoquinoline alkaloids have also been reported as anitfeedant against *Hyphantria cunea* and *Agelastica coerulea*, however, this was not the case in our study. We did not find ants refusing to eat this toothpaste extract, which further indicated its potential to be developed as bait agent. The results are similar to the toxicity of stone orchid powder to *Blattella germanica* Linnaeus, and the contact toxicity of new nicotinic insecticides to *Eurygaster integriceps* Puton. The toxicity effect was not significant when the concentration was low, but the toxicity was better at high concentration [20]. Apart from Yunhan Watermelon Cream Toothpaste, we also found that the toothpaste of Pianzaihuang Yahuoqing also had a high mortality to the red imported fire ants in the toxicity test. However, in the dyeing experiment, the living red fire ants treated with the toothpaste of Pianzaihuang Yahuoqing had a very low dyeing rate, which did not indicate whether the ants consume it or not. In this case, two assumptions can be made. One is that Pianzaihuang has high toxic activity to red fire ants, so the ants died quickly after eating it. Therefore, the living red fire ants cannot be found dyed. Secondly, the high mortality of red imported fire ants was caused by starving as they refuse to intake the toothpaste, may be due to repellency of toothpaste. Previous studies showed that *radix notoginseng*, (active ingredient of this tooth paste) contain large amount of ginsenosides, a triterpenoid saponin, which are strong repellant and deterrent to *Pieris rapae* and several other herbivore insect pests [8, 21].

Chinese medicine toothpaste contains traditional Chinese medicine, which are beneficial to oral health. Previous studies showed that the alcohol extract of traditional Chinese medicine has a good toxic effect against insect pests. For example, *Stemona japonica* can kill armyworm, aphid and *Tetranychus cinnabarinus*. The alcohol extract of *Celastrus angulatus, radix sophorae flavescentis, Siberia Cocklebur* and *Euphorbia kansui* had good contact poisoning on *Tetranychus cinnabarinus. Stellera chamaejasme, Stemona japonica, Veratrum nigrum*, and *Siberia Cocklebur* have a good contact poisoning on the cabbage aphid [22]. The traditional Chinese medicine contains a large amount of active substances, which can kill pests with low residue, and will not pollute the environment. The toxicity of the bait made by plant is slow, which is beneficial to spread in the red fire ant colony [23].

Therefore, in future we are focusing to test watermelon cream extracts under the field conditions and to figure out its active ingredients to develop effective insecticide’s formulations against this invasive insect pest.

## Funding

This study was supported by the National Key Research and Development Project (2016YC1201200). The funders had no role in the study design, data collection and analysis, decision to publish, or preparation of the manuscript.

## Compliance with ethical standards

### Conflict of interests

The authors declare that there is no conflict of interests regarding this work.

### Ethical approval

Guidelines on ethical issues and international agreements were considered and complied with.

## Supporting information

**S1 Fig. S1** Extraction steps of Yunhan watermelon cream toothpaste.

**S2 Table 1.** List of toothpastes with their active ingredients, other components and name of manufacturer that were used in the experiments

